# Regenerative hallmarks of aging: Insights through the lens of *Pleurodeles waltl*

**DOI:** 10.1101/2022.09.13.507508

**Authors:** Georgios Tsissios, Gabriella Theodoroudis-Rapp, Weihao Chen, Anthony Sallese, Byran Smucker, Lake Ernst, Junfan Chen, Yiqi Xu, Sophia Ratvasky, Hui Wang, Katia Del Rio-Tsonis

## Abstract

**Background:** Aging and regeneration are heavily linked processes. While it is generally accepted that regenerative capacity declines with age, some vertebrates, such as newts, can bypass the deleterious effects of aging and successfully regenerate a lens throughout their lifetime.

**Results:** Here, we used Optical Coherence Tomography (OCT) to monitor the lens regeneration process of larvae, juvenile, and adult newts. While all three life stages were able to regenerate a lens through transdifferentiation of the dorsal iris pigment epithelial cells (iPECs), an age-related decline in the kinetics of the regeneration process was observed. Consistent with these findings, iPECs from older animals exhibited a delay in cell cycle re-entry. Furthermore, it was observed that clearance of the extracellular matrix (ECM) was delayed in older organisms.

**Conclusions:** Collectively, our results suggest that although lens regeneration capacity does not decline throughout the lifespan of newts, the intrinsic and extrinsic cellular changes caused by aging alter the kinetics of this process. By understanding how aging affects lens regeneration in newts, we can gain important insights for restoring the age-related regeneration decline observed in most vertebrates.

## 1. Introduction

Exposure to endogenous and exogenous stimuli, such as oxidative, genotoxic, ribosomal, and mitotic stress can have catastrophic effects on cell state and function (Gorgoulis et al., 2019). Even though defense mechanisms like DNA damage response (DDR) and reactive oxygen species (ROS) management function to restore cellular harmony and maintain tissue homeostasis, the continuous exposure to stressors due to aging often leads to irreversible cellular alterations (Schumacher, 2009; Warraich et al., 2020; Yousefzadeh et al., 2021). Aging can cause a myriad of intrinsic and extrinsic cellular changes, including but not limited to, mitochondrial dysfunction, telomere attrition, epigenetic modifications, proteostasis impairment, as well as ECM and cell adhesion alterations (López-Otín et al., 2013; Sharpless and DePinho, 2007; Xiao et al., 2015). Consequently, these cellular alterations lead to physiological imbalance, immune system dysfunction, tissue and organ degeneration, disease vulnerability, and decline in regeneration capacity (Ring et al., 2022).

Humans cannot yet escape these age-related consequences (Gardiner, 2005; Xiao et al., 2015). However, regenerative competent species, like newts, show negligible signs of aging and retain tremendous regenerative abilities even at an older age (Arenas Gómez and Echeverri, 2021; Brockes and Kumar, 2008; Joven et al., 2019; Sánchez Alvarado and Tsonis, 2006; Sousounis et al., 2014b). In fact, to test the extent to which newts can regenerate, a series of truly astonishing studies were conducted, where newt lenses were surgically removed up to19 times over the period of 18 years from the same animals. By the time the final lenses were removed, the animals were 32 years old, and yet still regenerated perfect lenses (Eguchi et al., 2011; Sousounis et al., 2015). When compared at the transcriptomic level, no significant differences were observed between the lenses collected from the animals that have undergone lens regeneration 19 times and the lenses from animals that have never undergone regeneration (Sousounis et al., 2015). Interestingly, when tails from the same animals that had never undergone regeneration were compared with tails collected from younger animals, thousands of genes associated with aging were found to be differentially regulated. Most noticeable, electron transport chain-associated genes were found to be downregulated in older tails, an indicative sign of mitochondrial dysfunction and metabolic alterations.

It is worth noting here that axolotls and frogs can also regenerate their lens but lose this ability soon after embryonic development and metamorphosis respectively (Henry and Tsonis, 2010; Suetsugu-Maki et al., 2012; Vergara et al., 2018). When the iris of young axolotls, which serves as the source for lens regeneration, was compared at the molecular level with the iris of older regeneration incompetent axolotls, signs of aging were evident. Similar to newt tails, genes associated with electron transport chain, cell cycle, and DNA repair were significantly downregulated in the iris of old axolotls compared to young regeneration-competent animals (Sousounis et al., 2014a).

Based on the aforementioned studies, it is tempting to speculate that the initiation of regeneration triggers the rejuvenation of tissues and the surrounding microenvironment. However, the mechanism by which regenerative-competent animals avoid aging hallmarks to proceed with regeneration remains unclear. Understanding how aging affects cells at the cellular and extracellular levels in regeneration-competent animals is vital in our quest towards developing treatments for regenerative medicine and age-related pathologies. This is the first study to delineate the differences in cellular mechanisms and kinetics on lens regeneration across three developmental stages of newts: pre-metamorphic larvae, post-metamorphic juvenile, and adult. Using live imaging (OCT), cell proliferation assays (EdU labeling) and ECM staining we investigated the early stages of lens regeneration, in which iPECs reprogram to switch cell fate and become lens cells. We then looked at the later stages where the lens vesicle develops into a mature lens using 3D OCT imaging/processing and a statistical model to predict lens volume.

## 2. Materials and methods

### 2.1. Animal Handling and Ethical Statement

All experiments described in this study were executed under the guidelines provided by the Institutional Animal Care and Use Committee (IACUC) at Miami University. *Pleurodeles waltl* newts were housed in a homemade aquatic circulating system and daily care was performed following previously established husbandry guidelines (Joven et al., 2015). Environmental conditions such as temperature, feeding frequency, light cycle, water parameters and cleaning procedures were kept the same for all three age groups.

### 2.2 Surgical Procedures

Surgical removal of lens (lentectomy) and eye enucleation surgeries were performed as described before (Tsissios et al., 2022). Briefly, using a scalpel, a cut was made through the cornea and the lens was carefully removed with fine tweezers. For enucleations, the entire eyeball was removed by cutting the muscles and membranes around the sclera. The three age groups of *Pleurodeles waltl* consisted of: 2 month old pre-metamorphic larvae, 6 month old post-metamorphic juveniles, and 3 year old post-metamorphic adults.

### 2.3 Specatral-Domain Optical Coherence Tomography

A customized Spectral-Domain Optical Coherence Tomography (SD-OCT) was used to follow the lens regeneration process from the same animals. For a detailed description of the SD-OCT setup, refer to Chen et al. (Chen et al., 2021). In brief, a broadband light source centered at 850 nm was used in the SD-OCT, which can provide ~5 μm axial resolution and 7.5 μm lateral resolution respectively. A total of 2048 A-Scans were captured in each B-Scan, and 500 orthogonal positions across the lens were scanned to reconstruct a 3D image. Prior to imaging, newts were anesthetized and placed on a customized stage to avoid involuntary motion.

### 2.4 Live Imaging and Three-Dimensional Reconstructions

A total of 6 animals (12 eyes) were followed by OCT for 356 days. At first, images were taken every other day from day 0 until 28 days post-lentectomy (dpl) in order to capture all the dynamic changes that occur during the early stages of lens regeneration. Afterward, OCT imaging was performed every 5 days up until 88 dpl, and then every 50 days up until 365 dpl. B-scan images were cropped and rescaled to the accurate ratio according to the scanning distance of each individual eye. Then, the C-Scan stacks were reconstructed with B-Scans, and the lens tissue was segmented separately to synthesize the color label depicted in Fig. 2.

### 2.5 EdU Injections, Histology, and Cytochemistry

Newt eyes were collected at 1,4,10, and 15 dpl for histological examination. At 24 hours prior to collection, EdU (Invitrogen, #C10338) was added intraperitoneal at 10 ug/g of BW. Following eye enucleation, the entire eyeball was fixed in 10% formalin overnight at 4 *°C* and paraffin embedding was performed as described previously (Tsissios et al., 2022). Histological sections of 10 μm thickness were obtained and EdU and picrosirius red staining (Polysciences, #24901) were performed according to the manufacturer’s protocol.

### 2.6 EdU Quantification and Statistical Analysis

The cell cycle re-entry experiment included the effects of two factors on the ratio of EdU+ to Hoechst stained cells: age (larvae, juvenile, adult) and time (1,4, 10, and 15 dpl). For each of the twelve combinations of the two factors, six eyes were processed, leading to a total of 72 experimental units. Each of the experimental units was reasonably assumed to be independent. Cells from nine cross-sections for each eye were counted, and the total number of EdU+/ Hoechst cells in both the dorsal and ventral iris were used in the statistical analysis. Because these measurements are counts, the appropriate statistical analysis is based on the negative binomial distribution (Lindén and Mäntyniemi, 2011; Ver Hoef and Boveng, 2007; White and Bennetts, 1996). Thus, a negative binomial regression, with two factors (day and time) as well as the interaction between the two factors, was used to model the number of EdU+ cells. Additionally, the number of Hoechst cells was included as the offset, which allows us to model the ratio of EdU to Hoechst cells. Using the interaction model with offset, the adult ratio was compared to the larvae and juvenile ratios, at each time point, leading to a total of eight comparisons. The eight p-values were adjusted by controlling the False Discovery Rate (Benjamini and Hochberg, 1995).

### 2.7 Lens Volume Quantification and Statistical Analysis

The volume of the lens was measured with SD-OCT as described before (Chen et al., 2021). The same SD-OCT images that were obtained from section 2.4 were used to calculate the volume. Each eye is assumed to be independent of every other eye, with two animals and thus four eyes in each age category (larvae, juvenile, and adult). Each eye was imaged every other day from 0 to 28 dpl, as well as on day 33 and day 48, for a total of 14 measurements. One of the larvae animals died at 26 dpl, therefore lens volume after that time point was not included in the analysis. For each image, the volume of the lens was estimated, along with the diameter of the eye using the sectioning feature of ImageJ software. There are several aspects of this data that make it challenging to analyze. First, the trajectories are not easily modeled with standard polynomials. Second, the measurements over time on a particular eye cannot be assumed independent; they are likely correlated. Third, the variation in the data was smaller early in the regeneration process. To address the first problem, Generalized Additive Models were used, to provide great flexibility to model nonstandard mean structures (Wood, 2006; Yee and Mitchell, 1991). For the second issue, a Generalized Additive Mixed Model (GAMM) was fitted via the gamm function in the mgcv R package in order to account for the correlation between measurements on a particular eye across time (Wood, 2006; 2022). In particular, a continuous autoregressive error structure (corCAR1) was used which estimates the level of correlation between successive measurements and assumes that the correlation declines as the measurements get further away in time (Pinheiro and Bates, 2000). For the third complication, we modeled the natural log of the volume which stabilized the variance. The GAMM that we fit included smooth estimates of the mean log (Volume) for each level of age while accounting for the diameter variable in the model as well.

## 3. Results

### 3.1 Real-time monitoring of lens regeneration in adult, juvenile, and larvae newts

We used SD-OCT to follow the process of newt lens regeneration in the three different age groups. We first imaged the intact eyes. Most anteriorly of the newt eye lies the cornea, a multilayer tissue that protects the inner ocular tissues and allows light to pass through (Zhu et al., 2012). The amount of light the pupil lets through the lens is controlled by the iris which is composed of a stroma layer located on top of an iris epithelium. This stroma consists of heterogeneous cell populations including fibroblasts, endothelial cells, melanocytes, keratinocytes, mast cells, muscles, and blood vessels (Yamada and McDevitt, 1974). Below the stroma, the iPECs are fully packed with melanosomes and are aligned in a double layer to form the iris epithelium (Zhu et al., 2012). Due to the heavy pigmentation, iPECs, just as cells of the retinal pigment epithelium, exhibit high scattering properties, thus resulting in the brightest signal observed via SD-OCT (Meleppat et al., 2019; Wilk et al., 2017). Within the lens, the lens epithelium and fibers are also distinguishable via our SD-OCT setup. Lastly, the space between the lens and the cornea (the anterior chamber) is filled with a transparent liquid known as the aqueous humor (Zhu et al., 2012) that appears clear via SD-OCT (Fig. 1 IV-VI). Before lens removal, apart from the obvious size variations, no morphological or gross-anatomical differences were observed in the eyes between the three age groups with SD-OCT imaging (Fig. 1 IV-VI).

**Fig. 1:**
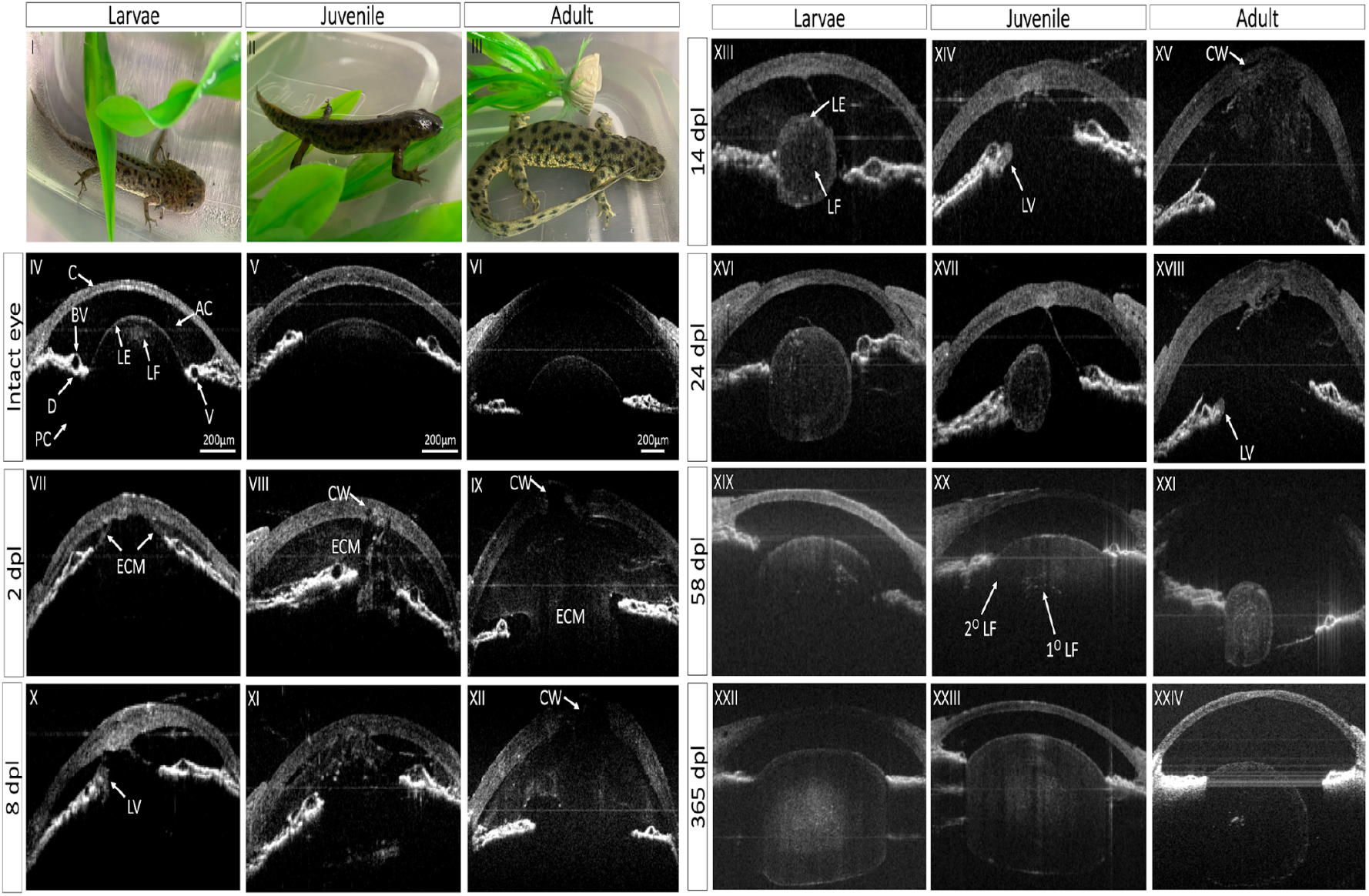
*In vivo* monitoring of lens regeneration from larval, juvenile, and adult newts. **A:** B-scan images are shown here in greyscale images. The eye was kept at the same orientation for all time points, with cornea (C) at the anterior, dorsal iris (D) on the left side, and ventral iris (V) to the right of the image. This figure depicts the timing of morphological events that occur during lens regeneration in each age group such as healing of cornea wound (CW), extracellular matrix (ECM) remodeling in the anterior (AC) and posterior chambers (PC), lens vesicle (LV) appearance, primary lens fiber (1°LF) differentiation from the epithelial cells at the internal layer (IL) of the lens vesicle, and lens epithelial differentiation into secondary lens fibers (2°LF).

**Fig. 2:**
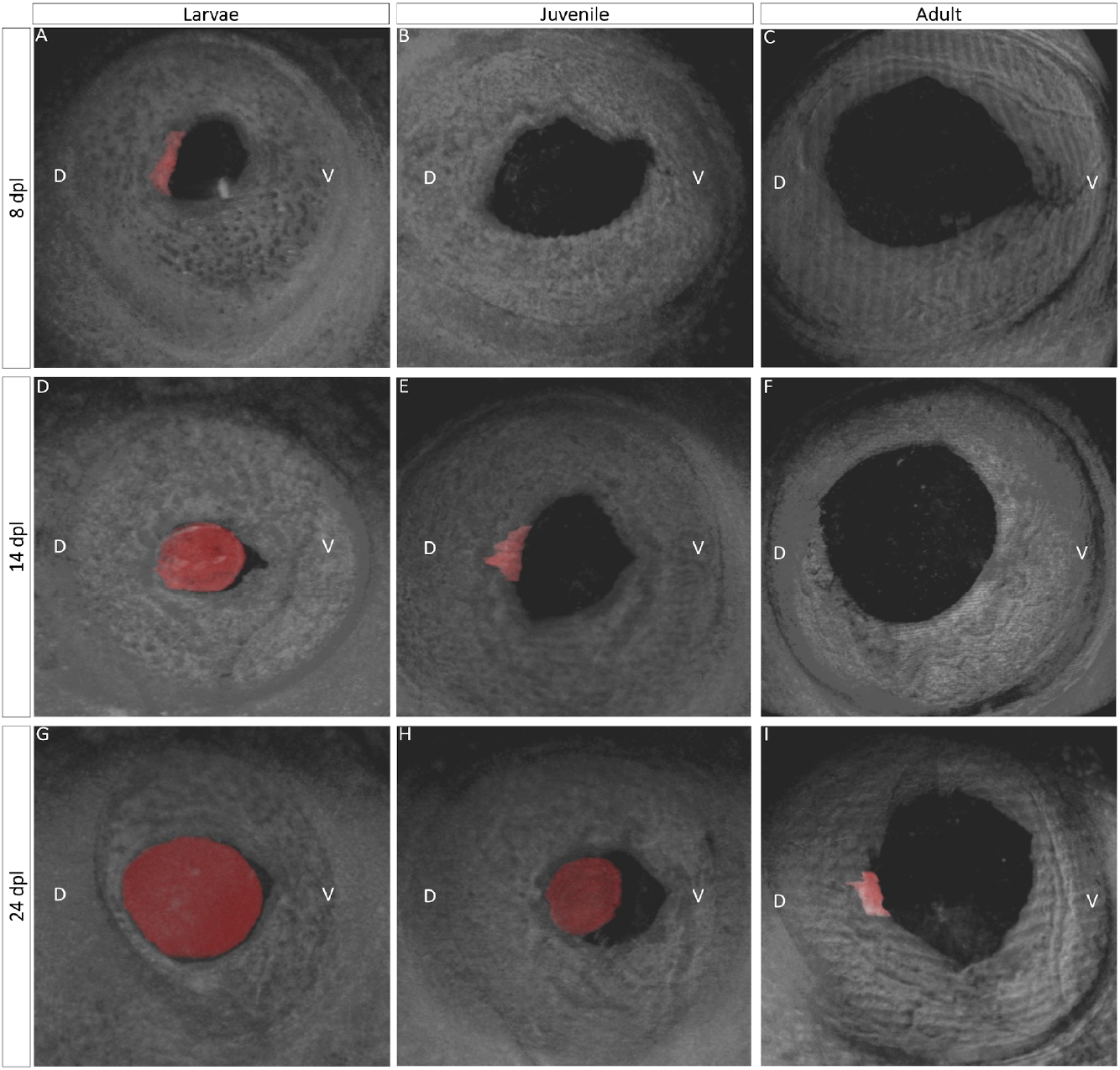
Three-dimensional view of lens regeneration. B-scans were used to reconstruct a threedimensional image of the newt eye. For easier visualization, the regenerating lens was pseudo-colored with red color. Three-dimensional images permit detailed observations of lens growth and morphogenesis. When it first appears, the lens vesicle is located in the mid-dorsal region of the iris and has an asymmetrical shape (A, E, I). As the lens develops and more lens epithelial cells differentiate into lens fibers, the regenerating lens assumes a spherical shape (D, G, H).

Lentectomy causes the loss of the aqueous humor and disruption between the anterior and posterior eye chambers. As a result, the anterior chamber is filled with ECM. Interestingly at 2 dpl, less ECM was observed in the anterior chamber of the larvae compared to juvenile and adult animals (Fig. 1 VII-IX). At this time point, the cornea appeared thicker in all three age groups compared to the corresponding cornea of the intact eye, indicating that proliferation to close the wound has started. By 4 dpl, the cornea appeared healed in larvae but not in juvenile and adult animals (Supplementary Fig. 1 I-III). At 6 dpl, the ECM in the anterior chamber of the larvae animals was mostly cleared out and a new transparent chamber was established (Supplementary Fig. 1 IV-VI). On day 8, iPECs, located at the tip of the dorsal iris of larvae animals, have depigmented and switched cell fate to become lens cells. These cells then formed an immature, asymmetrical lens vesicle (Fig. 1 X). By 12 dpl, the cornea of the juvenile has healed, and a clear anterior chamber devoid of ECM was observed (Supplementary Fig. 1 XI). At this time, ECM degradation is apparent in adult animals (Supplementary Fig. 1 XII). Later, at 14 dpl, a lens vesicle was visible in the juvenile (Fig. 1 XIV). As lens morphogenesis proceeds in larvae animals, elongated lens cells fill the lumen of the lens. These cells are called primary lens fibers and are arranged in a concentric fashion in the middle of the lens, whereas the cuboidal lens epithelial cells are arranged in a monolayer at the anterior periphery of the lens (Fig. 1 XIII). In adults, ECM did not completely clear until 22 dpl, and a lens vesicle appeared soon after at 24 dpl (Supplementary Fig. 1 XXIV, Fig. 1 XVIII).

During further growth, lens epithelial cells at the equatorial region of the lens differentiate and give rise to secondary lens fibers that surround the previously formed primary fibers. These newly formed lens fibers have different organelle compositions compared to primary lens fibers that have lost all organelles. The difference in organelles composition results in different scattering properties and therefore can be distinguished by SD-OCT (Fig. 1 XX; Supplementary Fig. 1 XXV, XXXVI) (Gupta et al., 2020). Eventually, the regenerating lens buds off from the dorsal iris to regain its original position. This is an essential step for regaining functional vision and a signal to mark the end of the regeneration process. In larval animals, the lens appeared detached from the dorsal iris at 33 dpl, and 58 dpl in juvenile animals (Supplementary Fig. 1 XXXI, Fig. 1 XX). Surprisingly, even at 365 dpl, the adult lens was still attached to the dorsal iris (Fig. 1 XXIV).

### 3.2 Three-dimensional view of lens morphogenesis

Three-dimensional SD-OCT images compiled from cross-sectional B-scans were used to acquire a multi-angle, panoramic view of the entire anterior eye. The first appearance of a lens structure (shown in red) was evident at 8 dpl from the mid-dorsal margin of the iris epithelium of larval animals, whereas the lens structure was noticeable at 14 dpl and 24 dpl in juveniles and adults (Fig. 2A, E, I). As the iris to lentoid transition takes place, the newly formed lens vesicle appears in an irregular, asymmetric and hollow shape. By 14 dpl in the larvae and 24 dpl in the juvenile, lens epithelial cells at the equatorial region of the regenerating lens elongate and differentiate into lens fibers, resulting in a larger and more symmetric lens (Fig. 2 D, H, Suppl. movie 1). As the regenerating lens continues to develop, newly formed secondary lens fibers symmetrically enclose the primary lens fibers, and the lens appears larger and more spherical at 24 dpl in the larvae. (Fig. 2G, Suppl. movie 1).

### 3.3 Growth rates of the regenerating lens

SD-OCT was used to quantify lens volume, and mean growth rates of the regenerating lenses were estimated using a Generalized Additive Mixed Model. The experiment makes it clear that the time by which the lens begins to grow is associated with age (Fig. 3A). Once the growth begins, however, the trajectories of juvenile and adult were very similar. However, there is evidence that the larvae trajectory for lens growth is different from both the juvenile and adult (Fig. 3B; p < 0.001) when comparing the larvae to the other two smoothed curves. For example, when measuring from the time point when growth was detected, the larvae started off smaller but quickly accelerated past juveniles and adults in terms of size.

**Fig. 3:**
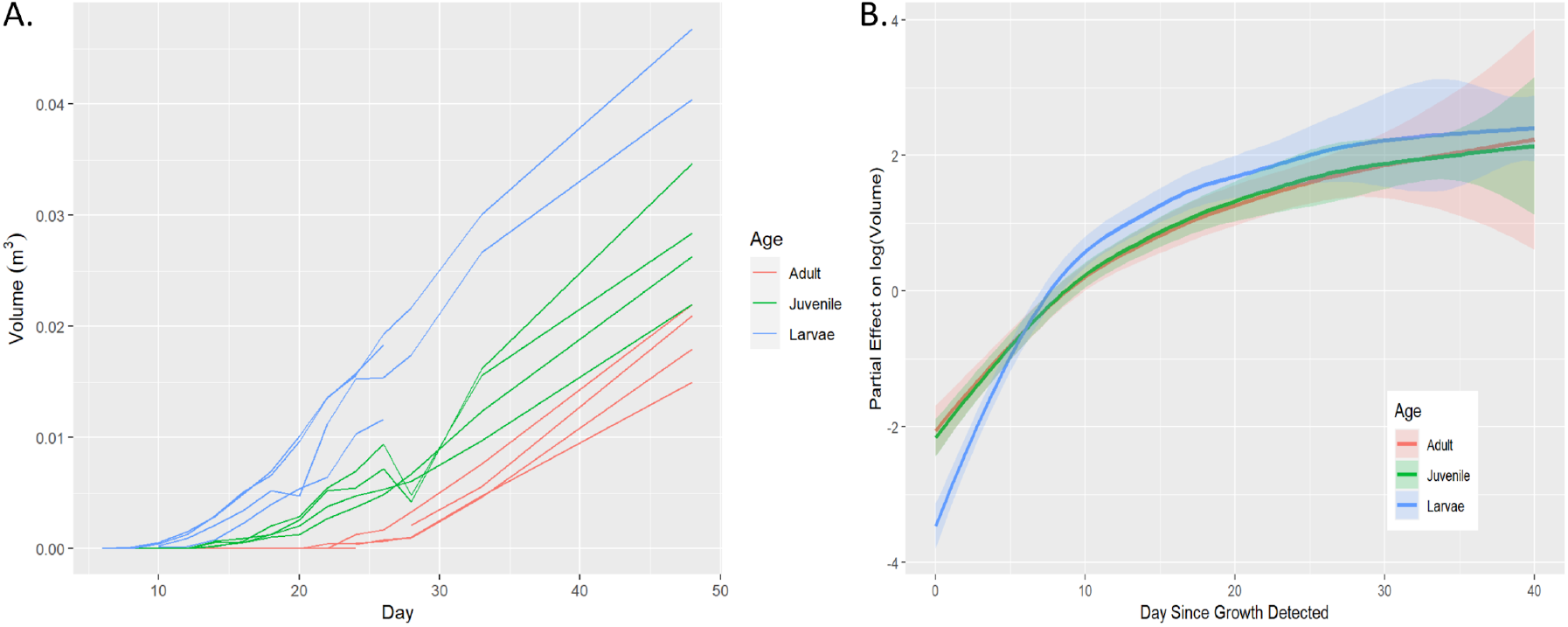
Comparisons of lens growth rates between three age groups. **A:** SD-OCT images were used to calculate the volume of lens vesicles from the time they appeared until 48 dpl. The raw volume data is plotted across the day of the experiment. **B:** A Generalized Additive Mixed Model (GAMM) was used to estimate the effect of the three groups on lens log (Volume) once the lens vesicle appears.

### 3.4 iPECs cell cycle kinetics

EdU+ cells were observed in the iris epithelium and stroma of larval animals as early as 1 dpl (Fig. 4A, B). At 4 dpl, cell cycle entry was evident in juvenile newts but at a much higher rate in larval animals. Once the lens formed, in larvae, iPECs that did not transdifferentiate to become lens cells withdrew from the cell cycle, and eventually returned to near baseline levels at 15 dpl (Fig. 4A, B). A significant delay in the iPECs cell cycle entry was observed in the adult eye, and no signs of fully depigmented iPECs were noticeable during the first 15 days.

**Fig. 4:**
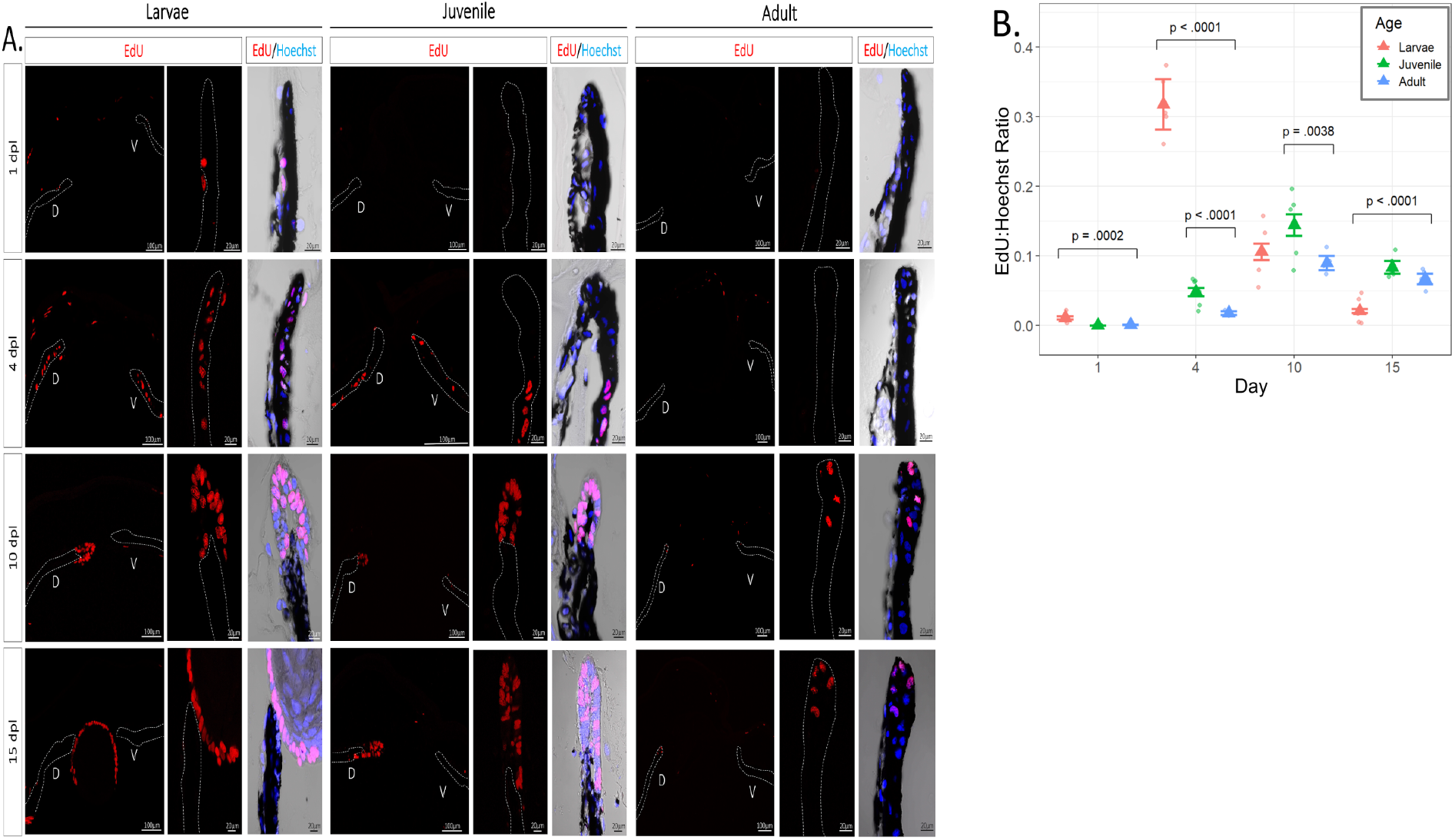
Histological examination and quantification of cell cycle dynamics during lens regeneration. **A:** Click-it EdU staining was performed to explore differences in cell cycle re-entrance and progression between the three age groups. EdU cells were observed in the iris epithelium (dotted line) and stroma of larval eyes at 1 dpl and in higher numbers at 4 dpl. By 10 dpl once the lens vesicle is formed no EdU cells were observed in the iris epithelium. EdU cells were found at 4 dpl in juvenile eyes and not until 10 dpl in adult eyes. **B:** The ratio of the number of iris epithelium EdUcells to Hoechstcells was plotted along the y-axis and the day lentectomy on the x-axis. Raw data (points) estimated mean ratio (triangle), and error bars using the standard error of the estimates are all displayed in the graph. Among the eight within-day comparisons with adult eyes, those with small FDR-adjusted p-values are annotated; none of the other three had p-values less than 0.2.

### 3.5 Collagen synthesis and remodeling during lens regeneration

Collagen staining was performed to confirm SD-OCT observations about ECM remodeling during regeneration. At 1 dpl, juvenile and adult eyes showed strong picrosirius red staining in the anterior and posterior chambers, indicating a homogenous presence of dense collagen fibers and sheets in the entire eye (Fig. 5). However, this was not the case in larval newts, as non-homogeneous collagen staining was observed throughout the eye cavity. In the anterior chamber, collagen appeared compacted, whereas collagen presence was less dense in the rest of the eye (Fig. 5). Collagen remodeling in the larvae eyes continued at 4 dpl, and by 10 dpl, most collagen was cleared out from the anterior chamber. Juvenile newts depicted clearing of collagen at 15 dpl, whereas in adult animals, the aqueous chamber contained a high percentage of collagen even at 15 dpl when the first signs of collagen remodeling were observed.

**Fig. 5:**
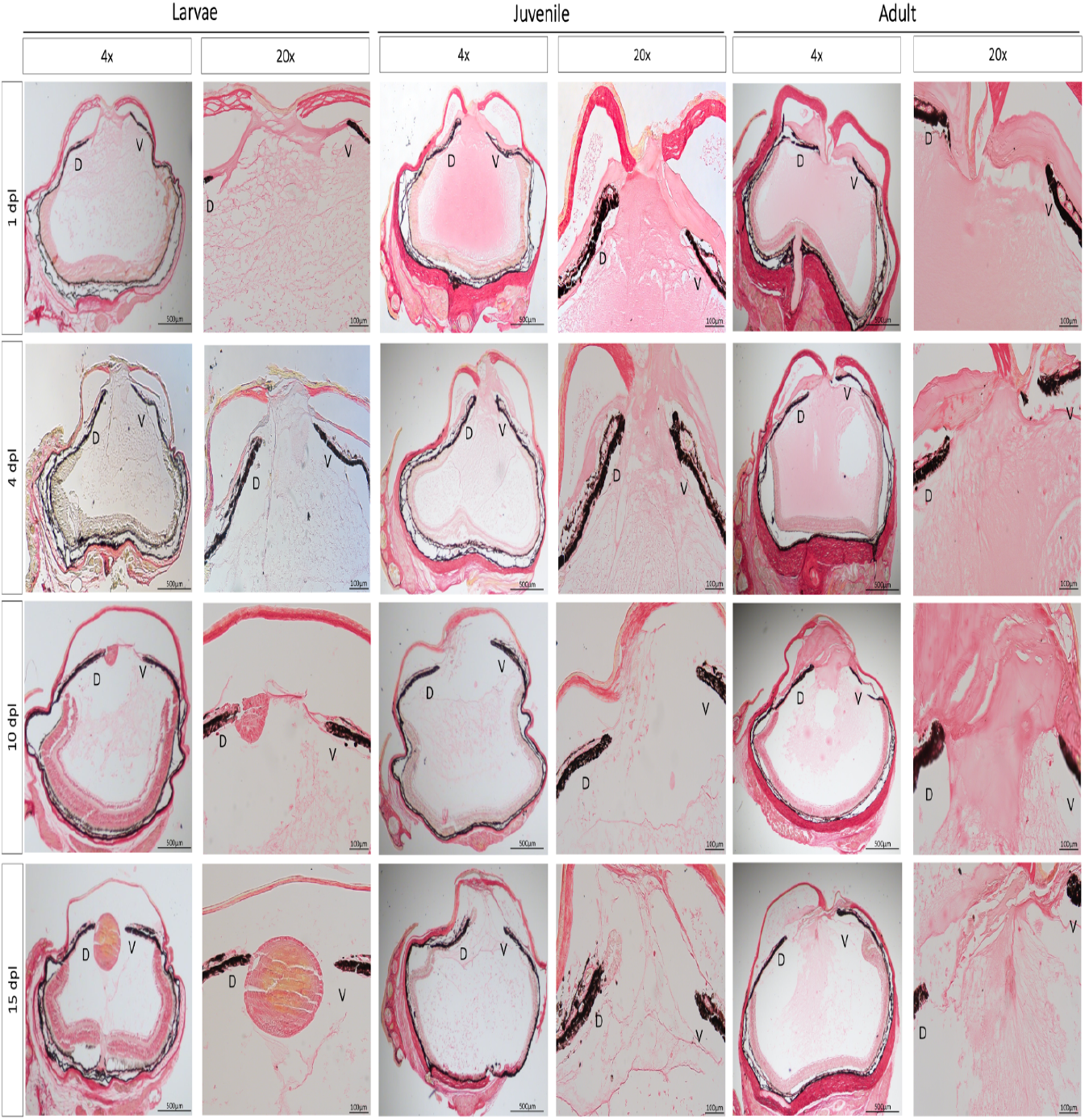
Histological examination of ECM remodeling during the early stages of lens regeneration. Picrosirius red staining was performed to visualize collagen composition and distribution throughout the newt eye at different stages of lens regeneration. Breakdown of collagen fibers in the anterior chamber is evident as early as 1 dpl in larval animals and by 10 dpl almost all collagen is cleared out of the eye chamber. On the contrary, strong collagen staining was observed all over the eye cavity in juvenile and adult eyes at 1 dpl. The remodeling process occurs slower in adult animals, as evidenced by the collagen staining detected in the anterior chamber at 15 dpl. D= dorsal iris; V= ventral iris.

## 4. Discussion

In this study, we demonstrated that although the capacity for newt lens regeneration is not lost with aging, the dynamics of this process are critically altered. Using *in vivo* imaging, we were able to follow the process of lens regeneration uninterrupted for 365 days from the same individuals and reported the morphological and cellular changes that occurred during the various stages of this process. We found that during the early stages of transdifferentiation, phenomena such as cornea healing, ECM remodeling, iPECs cell cycle re-entry, and lens vesicle appearance occurred faster in the pre-metamorphotic larvae compared to post-metamorphotic juvenile and adult animals. On the other hand, no clear differences were observed in the lens development rate and morphogenesis during the late stages of lens regeneration.

Although the cellular events that govern lens regeneration have been previously described in newt species, such as *Notophthalmus viridescens* and *Cynops pyrrhogaster*, to our knowledge this is the first detailed report that characterized stages of lens regeneration in *Pleurodeles waltl*(Eguchi, 1963, 1964; Reyer, 1954; Vergara et al., 2018; Yamada, 1977). Tadao Sato categorized the process of lens regeneration into 13 distinct stages (Sato, 1930; 1935). In the early stages, iPECs undergo transdifferentiation to reprogram and become lens cells (Sato stages 1-3). We observed a drastic difference in the timing of lens vesicle appearance among the three age groups. A lens vesicle was detected as early as 8 dpl in larvae, 14 dpl in juvenile, and 24 dpl in adult animals. After a lens vesicle is formed, the mechanisms and events of lens growth and morphogenesis parallel those of eye development in vertebrates (Tsonis et al., 2004). At first, epithelial cells at the internal wall of the lens vesicle become elongated to form the primary lens fibers (Sato Stage 7-8). Sequentially, the epithelial cells at the equatorial region of the lens stop proliferating, begin to change shape, and then lose their organelles as they differentiate into lens fibers (Chaffee et al., 2014). These secondary lens fibers then migrate towards the center of the lens, where they will surround the previously formed primary fibers (Sato Stages 9-11). No obvious differences were observed regarding the aforementioned timeline, as well as the speed of lens development and morphogenesis. To further compare the rates of lens growth between larvae, juvenile and adult animals, three-dimensional SD-OCT images were utilized to quantify lens volume growth. This method has several advantages as it allows us to continuously quantify the lens growth from the same animal without any tissue artifacts that are often introduced via histological processing (Chen et al., 2021). Consistent with our morphological observations, after the lens vesicle was formed, we observed similar rates of growth between juvenile and adult lenses, whereas the size of the larval lens was initially smaller but grew more rapidly until about 10 dpl.

Once the lens is mature during the late stages of lens regeneration and covers the entire pupil area (from dorsal to ventral iris), it detaches from the dorsal iris and relocates to its initial position in the middle of the eye where it is held by the zonule fibers (Tsonis et al., 2004). Interestingly, even though the rate of lens growth was not different, we observed a delay in lens detachment from the dorsal iris in older animals. In fact, even at 365 dpl, the lenses of adult newts were still attached to the dorsal iris. As our data displays a similar growth rate between age groups, we concluded that the reason the adult animal failed to reach the size of an intact lens by 365 dpl was not in fact due to differences in the speed of lens growth, but due to the fact that the size of the lens in adults are much bigger than the lens of younger animals. Therefore, it will take the aged newts a longer time to obtain the necessary size. The smaller the original lens, the faster the regenerating tissue will reach its original size.

The re-entrance of iPECs into the cell cycle is the most prominent and critical step during the early stages of lens regeneration. Under normal conditions, iPECs are terminally differentiated cells that show no signs of cell division (Eguchi and Shingai, 1971; Reyer, 1971; Yamada and Roesel, 1969; Zalik and Yamada, 1967). Lentectomy reverts the differentiation status of these mature cells and triggers cell cycle re-entry. Cell cycle kinetics during lens regeneration have been well documented both *in vivo* and *in vitro* in a variety of newt species (Horstman and Zalik, 1974; Yamada et al., 1975; Zalik and Yamada, 1967). From these studies, it was concluded that iPECs transition from G_0_ to G_1_ between 3-5 dpl and start to withdraw from the cell cycle once the lens vesicle is formed. Here we observed that cell cycle re-entry was faster and at a higher magnitude in larval animals. Surprisingly, we detected EdU-positive cells from the epithelium and stroma of the larval iris as early as 1 dpl. At 4 dpl, almost the entire dorsal and ventral iris was EdU positive. By 10 dpl, once the lens vesicle was formed, the EdU:Hoechst ratio was decreased as the iPECs withdrew from the cell cycle and regained their original identity. In contrast, very few EdU+ cells were observed in adult eyes at 10 and 15 dpl. Furthermore, no signs of depigmentation were observed until 15 dpl. It is interesting to note here, that when the process of lens regeneration was investigated in outer space, an environment known to cause anti-aging effects in humans, cell cycle re-entry was faster and at a higher magnitude when compared to newts that were kept on Earth (Grigoryan et al., 2002; Otsuka et al., 2021). Further exploring the association between microgravity, aging, and cell cycle acceleration could provide important insights into the aging effects of iPECs seen in this study.

The composition and remodeling of ECM is a fundamental aspect of wound healing and critical determinant of the regeneration outcome (Erickson and Echeverri, 2018). Failure to remodel ECM during the early steps of wound healing leads to scar formation, whereas the dynamic degradation of ECM proteins is often associated with regeneration success (Arenas Gómez et al., 2020; Huang et al., 2021; Satoh et al., 2011; Seifert et al., 2012; Vinarsky et al., 2005). Despite the importance of ECM remodeling, very little is known about this process during lens regeneration. Through histological examination and electron microscopy studies, it has been shown that ECM accumulates into the anterior aqueous chamber immediately after lens removal (Tsonis et al., 2004; Yamada, 1977). Prior to lentectomy, the anterior chamber of the eye is filled with a transparent liquid that allows light to pass through and support the function of lens epithelial cells. Following lentectomy, the transparent aqueous humor is lost, and the eye cavity fills with ECM. In this study, we used SD-OCT and picrosirius red staining to investigate the dynamic remodeling of ECM during regeneration and characterized age-related variations of this process. OCT imaging was previously validated as a powerful tool to detect real-time changes in ECM composition in several mammalian tissues (Carpaij et al., 2020; Yang et al., 2011). To the best of our knowledge, this is the first attempt to monitor ECM during lens regeneration *in vivo*. We observed across all three life stages that the lens vesicle did not develop until the extracellular matrix was completely cleared from the aqueous chamber. These results highlight the importance of ECM clearing for successful regeneration. A possible explanation for the effects of ECM on regeneration is that it interferes with the trafficking of growth factors produced by the retina. It was previously shown that growth factors such as FGFs are produced and secreted from the retina and that FGFs are necessary for lens regeneration in the newt eye (Caruelle et al., 1989; Del Rio-Tsonis et al., 1997; Del Rio-Tsonis et al., 1998; Hayashi et al., 2004). Therefore, we hypothesize that the presence of dense matrix proteins in the eye cavity can negatively impact the trafficking of growth factors secreted from the retina and used by the iPECs.

## 5. Conclusion

Collectively, we have shown that aging delays the process of transdifferentiation during lens regeneration in *Pleurodeles waltl*. Several theories can be drawn to explain the delay in transdifferentiation from older newts. One possible explanation is that the differentiation status of iPECs is more advanced in adult animals, causing the dedifferentiation process to take longer. Future work should explore the molecular signatures of iPECs from young and old animals during lens regeneration, which will provide age-related differences in differentiation trajectories and cell fate decisions.

An increased accumulation of senescent cells could also account for the delay in regeneration from older newts. Senescent cells secrete numerous growth factors, proteases, cytokines, chemokines, and matrix remodeling proteins that can negatively impact several stages of regeneration (Yu and Yun, 2020; Yun, 2021). The importance of clearing senescent cells in mammalian tissues was highlighted by Baker et al., where genetic targeting of senescent cells was shown to alleviate cellular dysfunction caused by aging (Baker et al., 2011). Interestingly, senescent cells were found to be present during limb regeneration in newts and axolotls but these are quickly recognized and removed by macrophages (Yun et al., 2015). Immune senescence could explain the delay in collagen clearing and cell cycle re-entry that we observed. Future studies will need to determine how aging affects macrophage function during regeneration and their ability to clear senescent cells.

Aging could also cause changes in pathways related to DNA damage response and genome stability that can negatively affect the process of transdifferentiation (Schumacher et al., 2021; Yousefzadeh et al., 2021). Microarray comparisons between iPECs from lens regeneration competent young axolotls and lens regeneration incompetent semi-adult axolotls revealed that the expression of several DNA repair genes is downregulated in older animals (Sousounis et al., 2014a). Furthermore, alteration of the DNA repair process in axolotls by genetic mutation and pharmacological inhibition of the gene Eya2 caused several limb regeneration defects and a delay in cell cycle dynamics (Sousounis et al., 2020). Future investigations are necessary to explore the influence of genotoxic stress on iPEC’s ability to reprogram into lens cells.

We recognize these theories are not mutually exclusive and the proposed mechanisms could be working together to promote the ontogenetic preservation of regenerative powers in newts. Unraveling the mechanisms by which regeneration-competent animals circumvent the deleterious effects of aging and retain tremendous regenerative powers will make major inroads in the development of therapies to treat geriatric diseases.

## Acknowledgments

The authors would like to thank Drs. Max Yun and Andras Simon from the Center for Regenerative Therapies in Dresden, Germany, and Karolinska Institute in Stockholm, Sweden for gifting us *Pleurodeles waltl* animals to start our colony at Miami University. Special thanks to Dr. Alberto Joven for his guidance in husbandry and breeding techniques. In addition, the authors thank Laboratory Animal Resources (LAR) director Jazzminn Hembree and the entire LAR team for their help to establish and maintain our colony. Moreover, further thanks to Matt Duley from the Center of Advanced Microscopy and Imaging (CAMI) at Miami University and Erika Grajales-Esquivel for their help and guidance on imaging procedures.

This research was supported by grants from the National Eye Institute: RO1 EY027801 (to KDRT) and R21 EY031865 (to HW and KDRT), by the John W. Steube Professorship Endowment (to KDRT), the Sigma Xi Aid in Research Grant (to GR) and Miami University undergraduate research grants.

