## Supplemental Figure 1 and movie 1 for "Regenerative hallmarks of aging: Insights through the lens of *Pleurodeles waltl*"

A. Supplementary data
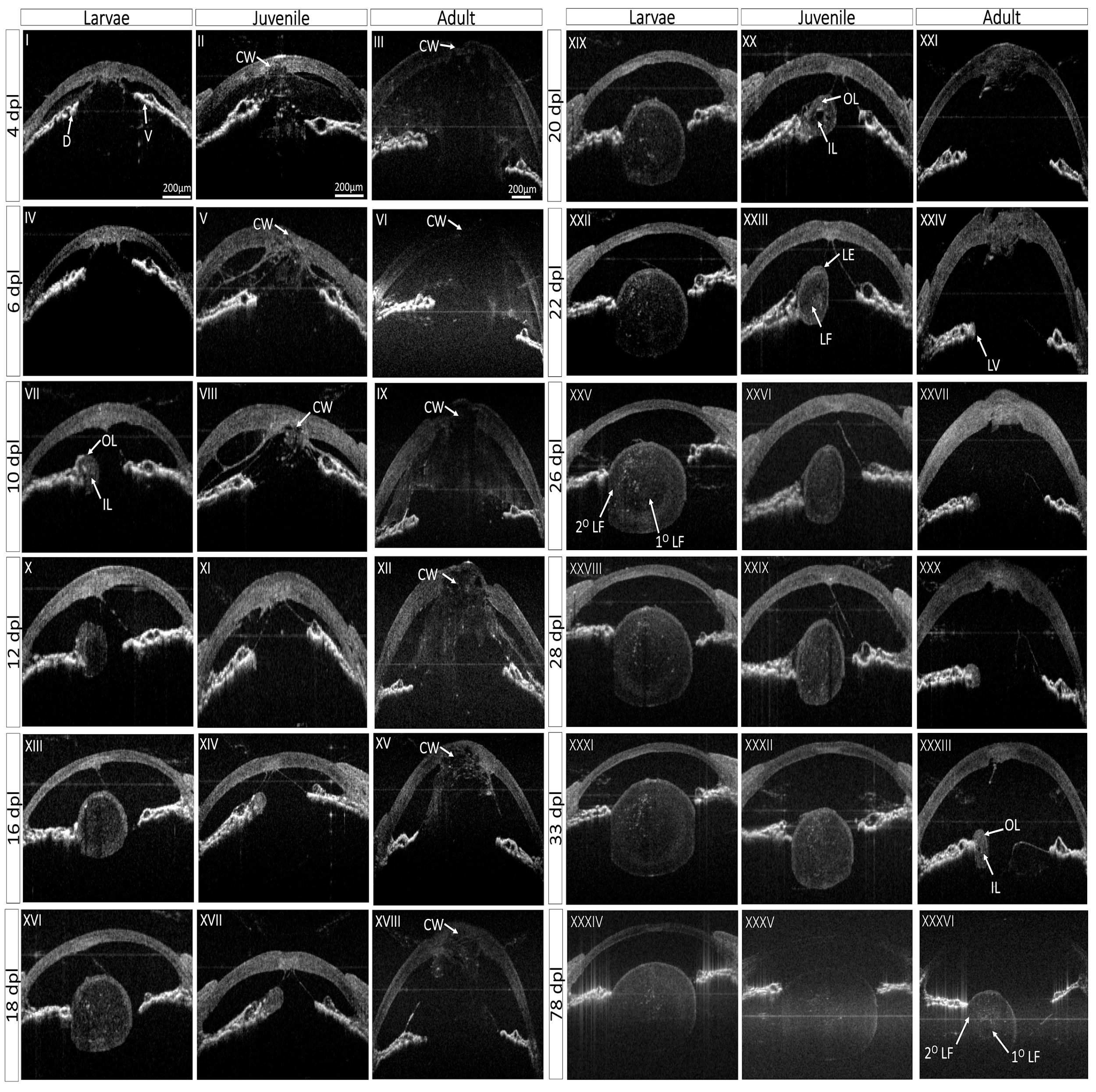

**Supplementary Fig. 1:** Additional time points of *in vivo* imaging across all three age groups. SD-OCT B-scans are shown here in greyscale images and the eye orientation has been kept the same throughout the figure. This figure depicts additional time points of the morphological events that occur during lens regeneration including cornea wound (CW), ECM remodeling, 1^o^LF and 2^o^LF differentiation, as well as the formation of the lens epithelium (LE) and lens fibers (LF).

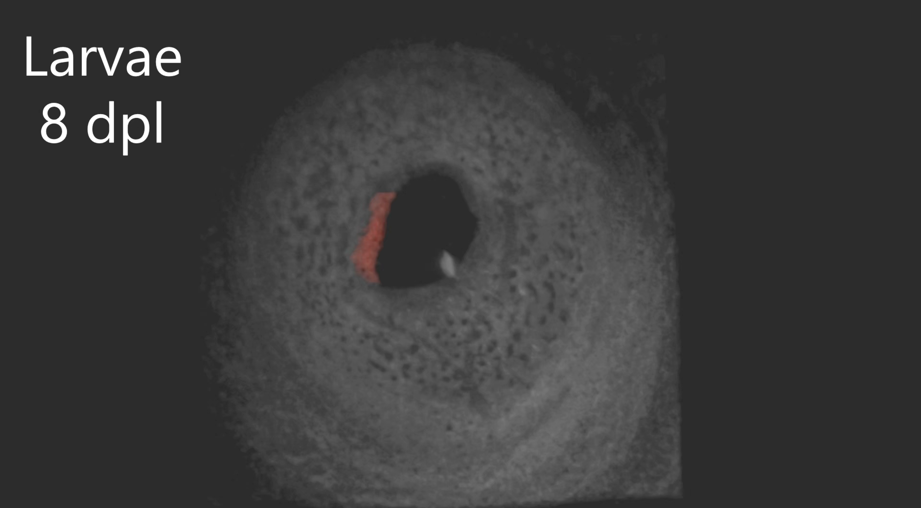

A.

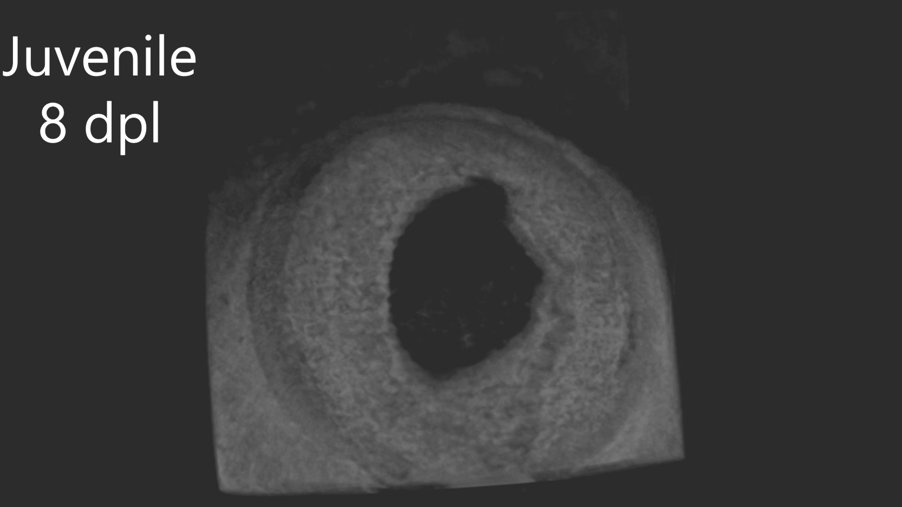

B.

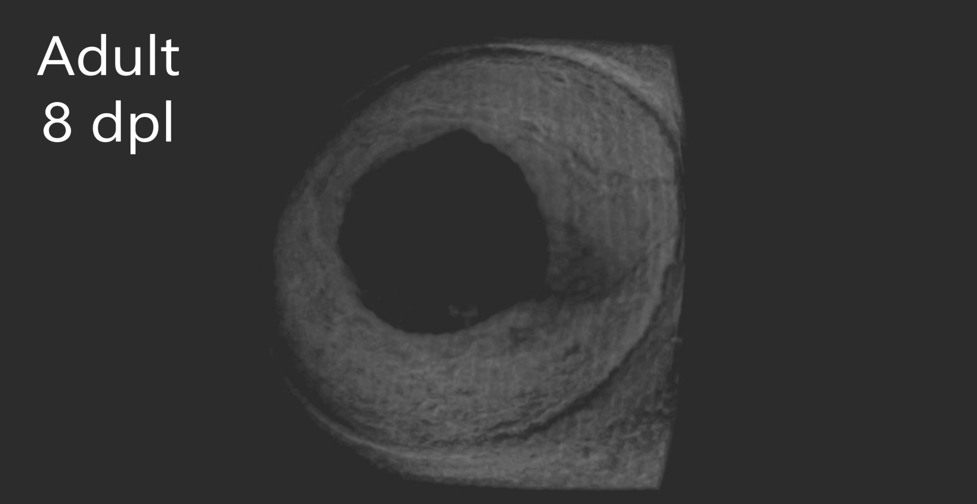

C.

**Supplementary Movie 1:** Rotating SD-OCT images of the anterior eye**.** The three-dimensional SD-OCT movie offers a spatial temporal view of the lens regeneration kinetics from larvae (A), juvenile (B) and adult (C). To aid in visualization, the lens was labeled in red color.
